# Low fidelity assembly of influenza A virus promotes escape from host cells

**DOI:** 10.1101/332650

**Authors:** Michael D. Vahey, Daniel A. Fletcher

**Author notes:** Department of Biomedical Engineering, Washington University in St. Louis, St Louis, MO.

## Abstract

Influenza viruses inhabit a wide range of host environments using a limited repertoire of protein components. Unlike viruses with stereotyped shapes, influenza produces virions with significant morphological variability even within clonal populations. Whether this tendency to form pleiomorphic virions is coupled to compositional heterogeneity and whether it affects replicative fitness remains unclear. Here we address these questions by developing live strains of influenza A virus amenable to rapid compositional characterization through quantitative, site-specific labeling of viral proteins. Using these strains, we find that influenza A produces virions with broad variations in size and composition from even single infected cells. The virus leverages this phenotypic variability to survive environmental challenges including temperature changes and anti-virals. Complimenting genetic adaptations that act over larger populations and longer times, this ‘low fidelity’ assembly of influenza A virus allows small populations to survive environments that fluctuate over individual replication cycles.

## Introduction

In order to replicate, viruses must successfully navigate complex and unpredictable environments both inside and outside the host. Genetic variability within virus populations is central to this, helping viruses evade host immunity (Dingens et al., 2017), withstand antiviral drugs (Hirsch et al., 2000), expand host range (Ma et al., 2016; de Vries et al., 2017), and adopt new routes of transmission (Imai et al., 2012; Sorrell et al., 2009). In RNA viruses, the basis for genetic variability is virus-encoded error-prone polymerases that enhance the rate of mutation several orders of magnitude above that of the host (Holland et al., 1982; Pauly et al., 2017; Sanjuan et al., 2010). Combined with the large number of progeny (∼10^3^-10^4^) produced from an individual infected cell (Chen et al., 2007; Stray and Air, 2001), this allows viruses to sample a large genetic space more rapidly than their hosts, helping them to evolve around adaptive and therapeutic defenses, and to benefit from cooperative interactions among co-infecting quasispecies (Belshe et al., 1988; Clavel and Hance, 2004; Vignuzzi et al., 2006). Recently, large pools of genetic diversity have also been shown to be instrumental in the evolution of drug resistance via epistasis, allowing permissive mutations to reside in a population that can stabilize subsequent mutations conferring resistance (Bloom et al., 2010; Pennings, 2012; Rong et al., 2010; Wang et al., 2002).

In addition to genetic variability, many RNA viruses share significant morphological variability, as revealed by electron microscopy (Battisti et al., 2012; Bharat et al., 2012; Calder et al., 2010). Influenza A virus (IAV), an enveloped respiratory virus with recurring pandemic potential, is found in a morphologically heterogeneous ‘filamentous’ form that predominates in human isolates, with virions having a uniform diameter of ∼80-100 nm but lengths ranging in size from ∼100nm to several microns (Badham and Rossman, 2016; Chu et al., 1949; Dadonaite et al., 2016). Although this morphological variability is known to be influenced by the sequence of the viral matrix protein (Burleigh et al., 2005; Elleman and Barclay, 2004), relatively little is known regarding how it may influence infectivity and transmission or whether virus composition is similarly variable. Unlike viruses with highly ordered capsids and envelopes such as flaviviruses (Sirohi et al., 2016), the glycoprotein spikes of IAV do not appear to be highly ordered (Harris et al., 2006; Wasilewski et al., 2012), suggesting that IAV may lack a specific mechanism of tightly regulating envelope composition in addition to particle size.

Heterogeneity in virus morphology and composition, which we refer to together as phenotypic variability, could influence IAV transmission in a number of ways. As the virus releases from an infected cell, transports through the host environment, and eventually binds to another cell to start the next round of replication, IAV must form sufficiently stable attachments to host cells to infect, while still being able to break those attachments to escape after budding. To accomplish these competing tasks, IAV encodes separate proteins to bind host receptors (hemagglutinin, or HA, which binds to sialic acid) and destroy these same receptors (neuraminidase, or NA, which cleaves sialic acid), both of which are located on the virus envelope. On the cell surface or in other environments rich with secreted forms of sialic acid (e.g. gel forming mucins), NA activity is necessary for efficient virus release and transmission (Cohen et al., 2013; Palese et al., 1974), while HA is essential for viral attachment and entry into cells. Consistent with their essential but competing activities, the functional balance between HA and NA has been found to drive compensatory evolutionary responses when this balance is perturbed (Wagner et al., 2002). Indeed, the most widely used anti-influenza interventions, neuraminidase inhibitors, target and attempt to disrupt this balance, and succeed in reducing the duration of symptoms during susceptible infections by ∼25 % (Nicholson et al., 2000). If sialic acid binding and cleaving activities varied at the level of individual viruses due to differences in HA and NA packaging, the efficacy of treatments specifically targeting either protein could be reduced. However, little is known about how the abundance of HA and NA on individual virions varies within an IAV population.

We sought to determine whether the variable – or ‘low fidelity’ – influenza A virus assembly process that produces heterogeneity in virion morphology could also cause heterogeneity in HA and NA composition and, if so, whether this phenotypic variability aids in the transmission of progeny virions in unpredictable growth environments. Although the adaptive benefits of phenotypic variability have been documented in many other systems (Balaban, 2004; Baldwin, 1896; Rego et al., 2017), measuring its effects in viruses has traditionally been challenging. This is largely due to a lack of available methods for measuring virus phenotype – in particular, size, shape, protein abundance, and protein spatial organization – at the level of individual virions.

Current methods for investigating IAV composition and morphology are limited in their ability to link phenotype with replicative fitness. Mass spectrometry has provided important insight into the average abundance of both viral and host proteins at the level of virus populations (Hutchinson et al., 2014; Shaw et al., 2008), as well as helped map the landscape of post-translational modifications to which IAV proteins are subjected (Hutchinson et al., 2012). Electron microscopy has identified organizational features of individual viruses, from the arrangement of genomic segments (Noda et al., 2012) to the distribution of proteins on the surface of the virus (Wasilewski et al., 2012). However, it remains challenging to measure protein abundance at the single virus level across an entire population. More challenging still is to do so in a way that does not compromise the function of the proteins being measured or the replicative fitness of the virus as a whole so that complete infection cycles can be studied. To address this challenge and to characterize the intrinsic heterogeneity in genetically uniform populations of IAV, we developed strains of the virus harboring small (∼10 amino acid) tags on each of the virus’s most abundant structural proteins – HA, NA, M1, M2, and NP. Coupling small molecule fluorophores to these tags provides a handle by which we can quantify the abundance of specific proteins inside live, infected cells, or in individual released virions.

Using these fluorescently-tagged strains, we find that IAV composition and morphology vary widely, with total abundance of HA and NA each covering a ∼100-fold range and the length of released virions ranging up to 20 µm within a viral population. Remarkably, the same variation in HA and NA incorporation found in virions released from multicellular monolayers is also found in virions produced by a single infected cell, indicating that low fidelity in the assembly process itself generates compositional variability, rather than differences among infected cells. Our data also show that HA and NA abundance and virus length can change significantly and reversibly over the course of a single replication cycle, and that composition is coupled to morphology, with longer virions more enriched in HA relative to NA than shorter virions. Critically, we find that the observed phenotypic variability serves as a determinant of viral escape from the cell surface in response to temperature changes and exposure to neuraminidase inhibitors, underscoring the functional significance of morphological and compositional heterogeneity. The ability of IAV to produce virions with broad stoichiometric distributions of its two competing surface proteins, HA and NA, without making a genetic commitment favoring either protein, provides influenza A virus with an evolutionary hedge in complex and unpredictable environments.

## Results

### Creating fluorescently-tagged IAV strains

Fluorescence microscopy offers several attractive features that could address the challenge of capturing protein abundance of individual viruses across a population while maintaining infectivity. Labeling proteins with specific fluorophores, either chemically (Sletten and Bertozzi, 2009) or genetically (Cranfill et al., 2016), provides high contrast and allows proteins to be quantified and localized with high precision in a biochemically complex background. Additionally, fluorescence measurements can be performed on non-fixed, live samples, enabling the observation of dynamic phenomena that are inaccessible using electron microscopy. These features come at the cost of engineering tags into the virus that can be disruptive to virus replication; to date, efforts to rescue viruses harboring fluorescent fusion proteins on viral proteins have been limited (Lakdawala et al., 2014), presumably due to the large size (∼25 kDa) of the genetically appended molecules. An alternative approach is to use site-specific labeling to insert small tags (∼5-10 amino acids) into particular viral proteins which can then be recognized by an enzyme that catalyzes the attachment of a dye molecule, or which can be bound directly by a dye molecule with high affinity. The small size of these dye molecules and their associated tags (∼1 kDa each) reduce the likelihood that they will inhibit protein function. Site-specific labeling of IAV proteins has recently been reported for HA, NA, and NS1 (Li et al., 2010; Popp et al., 2012); however, this approach has not yet been harnessed to simultaneously and orthogonally label multiple viral proteins in live strains with replicative fitness comparable to wild type laboratory strains, a feature that is needed to study changes in virus composition and length over multiple rounds of replication.

We sought to construct IAV strains that are amenable to simultaneous fluorescence imaging of multiple viral proteins with minimal disruption to protein function and virus replication (Figure 1A). Using site-specific labeling via a combination of enzymes and probes – SFP synthase (Yin et al., 2006), Sortase A (Theile et al., 2013), transglutaminase 2 (Lin and Ting, 2006), and microbial transglutaminase (Sugimura et al., 2008) – as well as the enzyme-free and membrane-permeable dye FlAsH (Griffin, 1998), we were able to rescue multiple strains harboring three orthogonal tags, and one strain with four (SrtA-HA, ybbR-NA, M48-M2 and FlAsH-NP; Figures S1 & S2, supplemental text). Cells infected with these strains shed high titers of virus that can be collected, labeled, and immobilized on coverslips for imaging (Figure 1B) and analyzed by SDS-PAGE (Figure S3), and a similar strategy can be used to label the infected cells themselves (Figure S4). Viruses released from infected cells and immobilized on functionalized coverslips show heterogeneity in size and composition (Figure 1C). Large morphological variability is a hallmark of clinical isolates of influenza A virus, and is also seen in our engineered strain expressing M1 from A/Udorn/1972 in an A/WSN/1933 background, which produces particles ranging from sub-diffraction limited foci to filamentous virions up to ∼20µm in length. Although larger viruses (>1µm) are a relatively minor percentage of the total population (∼10%), they comprise roughly one third of the total viral material that buds from cells during infection (Figure 1D). We find that both the insertion of site-specific tags in the viral genome and the attachment of fluorophores to these tags are minimally disruptive; viruses containing multiple site-specific tags replicate with kinetics matching the parental strains (Figure 1E), and virus labeled with fluorophores maintain high infectivity (>70% of an unlabeled control) (Figure 1F).

**Figure 1:**
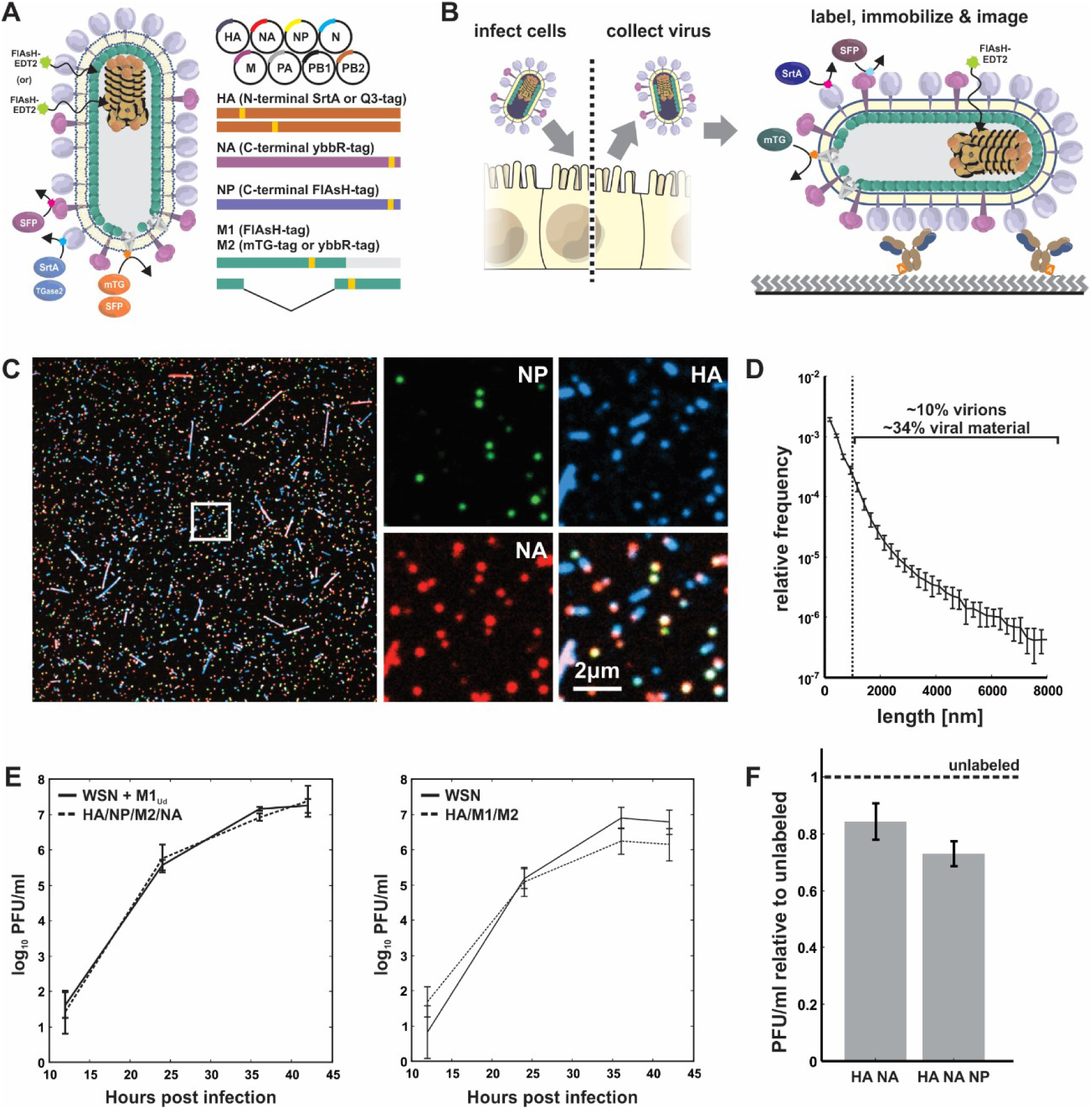
Multi-spectral strains of influenza A virus permit fluorescence-based quantification of virus composition and morphology. (A) Schematic of the virus labeling strategy. Inserting small (∼5-10aa) tags into the structural proteins of influenza A virus renders the virus amenable to labeling via the site-specific attachment of fluorophores. (B) Work flow for collecting and labeling IAV proteins on intact and infectious virions. (C) Viruses labeled and immobilized on a coverslip exhibit diverse morphology and composition. (D) Length distribution of virions containing M1 from A/Udorn/1972 showing significant size variation. Viruses >1μm in length comprise a minority of ∼10% of the population, but represent ∼34% of the viral material shed during infection (estimated as the product of measured particle size and frequency within the population). Plot shows mean and standard deviation from six biological replicates with N > 44000 viruses in each. (E) Viruses with up to four orthogonal tags show replication kinetics matching the parental strains in MDCK cells (error bars represent standard deviations of four independent replicates). (F) Labeling HA and NA on the surface of the virus preserves ∼85% infectivity relative to unlabeled tagged strains, while labeling HA, NA, and NP preserves ∼75% infectivity. Quantification shows the mean and standard deviation of labeled virus titers relative to unlabeled samples for three biological replicates.

At moderate multiplicity of infection (MOI ∼ 0.1-1), cells infected with tagged virus are clearly distinguishable from surrounding uninfected cells by the presence of fluorescence associated with each of the tags, allowing us to calculate the frequency with which pairs of viral genes are expressed in cells infected by ∼1 virus (Figure S5A). Cells labeling positive for any of the three membrane proteins (HA, NA, or M2) are positive when probed for a second membrane protein >90% of the time, and are positive for NP >99% of the time, consistent with the requirement for NP in the transcription of viral RNAs. Inversely, cells labeling positive for NP are positive for other viral proteins ∼75-85% of the time, a frequency which holds whether these proteins are labeled enzymatically or detected via an antibody that is independent of the engineered tags (Figure S5B & C). These results demonstrate the stability of the engineered tags within the viral population and are consistent with recent measurements suggesting that packaging of all eight segments of the IAV genome is up to 80% efficient (Nakatsu et al., 2016).

The abundance of viral proteins in particles immobilized on coverslips can be measured by quantifying fluorescence, which is carried out by thresholding intensity in the HA and NA channels to segment each particle and then integrating the intensity of fluorophores over the particle’s segmented area. In order to convert intensity values into absolute counts, we recorded images of labeled viruses using TIRF microscopy and calibrated the intensity of individual dye molecules under identical conditions. Correcting for the geometry of the virus and the shape of the evanescent field (supplemental text and Figure S6A & B) yields a median density of ∼6000 labeled HA molecules and ∼1100 labeled NA molecules per µm^2^ of virus area; for a spherical particle 120 nm in diameter, this corresponds to 517 HAs (∼170 trimers) and 93 NAs (∼23 tetramers). These values are comparable to those previously reported by mass spectrometry (294/98 HA monomers/trimers, 23/6 NA monomers/tetramers (Hutchinson et al., 2014)) and by direct counting of proteins on a similarly sized pair of viruses, imaged using cryoelectron tomography (870-903 HAs, 152-200 NAs (Harris et al., 2006)). Accounting for interactions between nearby dye molecules in our experiments would further increase the estimated HA and NA densities (supplemental text, Figure S6C & D).

### Low fidelity of influenza A virus assembly produces compositionally and morphologically diverse progeny

To determine if the protein composition of individual viruses is as heterogeneous as their morphology, we prepared viruses with HA, NA, or M2 labeled with Alexafluor 555, allowing us to measure the relative abundance of these proteins across populations of individual virions (Figure 2A). We find that HA, NA, and M2 abundance in individual virions each span a ∼100-fold range (with 50% of each population concentrated within a three- to five-fold range), reflecting significant compositional heterogeneity compared to viruses with stereotyped capsids and protein organization. This variation appears to be affected by morphology, as replacing M1 from A/Udorn/1972 with the sequence from A/WSN/1933 results in more monodisperse particles and a narrower distribution of HA abundance (Figure 2B). The relative medians of the distributions are in general agreement with published values from population measurements, with HA being approximately five-fold more abundant than NA and 40-fold more abundant than M2 (Hutchinson et al., 2014; Zebedee and Lamb, 1988). To compare the abundance of the internal viral proteins M1 and NP, we prepared separate virus samples in which one or the other of these proteins were labeled using FlAsH-EDT2. Owing to difficulties in rescuing viruses carrying FlAsH-tagged Udorn-M1 that preserve the filamentous morphology, these viruses contain WSN-M1 and produce predominantly spherical particles. Comparing the labeling intensities in these samples gives a median value for M1 that is roughly three-fold that of NP (Figure 2C), also consistent with previous measurements (Hutchinson et al., 2014; Oxford et al., 1981). As with the other viral proteins, there is considerable variation in M1 levels that could arise from differences in virus size, as well as particles lacking a complete matrix layer (Harris et al., 2006). The bimodal distribution of NP abundance we observe suggests that many of the virus-like particles that are released during infection (∼33%, based on fitting two log-normal distributions to the data) have missing or defective genomes. The percentage of NP-negative viruses is roughly twice as high in longer (>1µm) filamentous particles, suggesting a link between particle morphology and genome packaging (Figure S7A).

**Figure 2:**
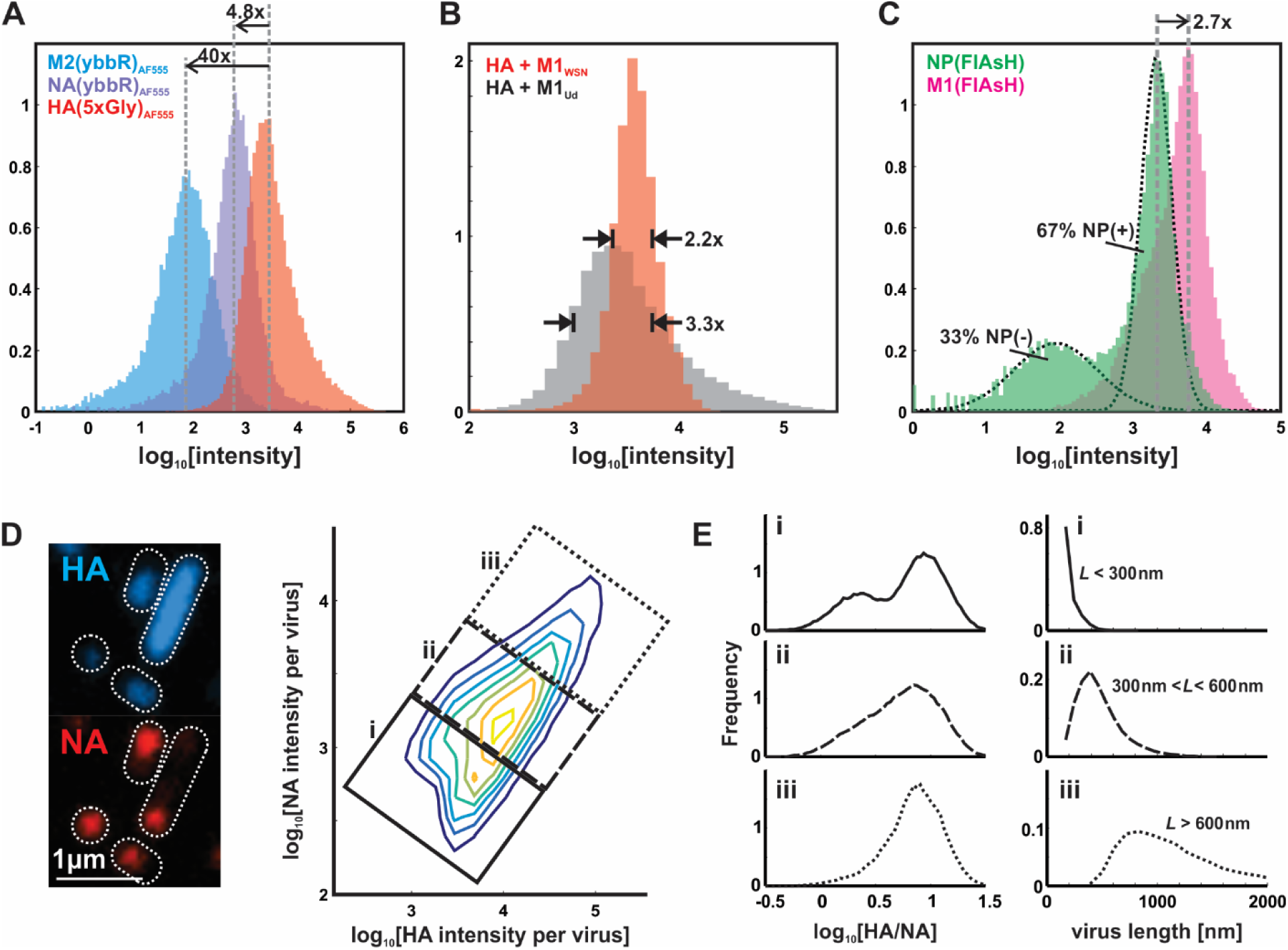
Influenza A virus composition and morphology vary broadly at the single-virion level. (A) Distributions of HA, NA, and M2 labeled via SFP synthase (NA and M2) and SrtA (HA). The relative medians of abundance are in agreement with prior measurements, but individual viruses vary considerably. This variation diminishes by ∼30% when M1_WSN_ is used to produce viruses of more homogeneous size. Similar distributions as in (A), but with NP and M1, quantified after labeling with FlAsH-EDT_2_. (D) Segmenting images of immobilized viruses and integrating the intensities of labeled HA and NA allows us to measure distributions of these proteins across a virus population, represented as a two-dimensional histogram showing the log-abundance of HA (horizontal axis) and NA (vertical axis) measured for 238419 viruses compiled from four independent experiments. Contrast in the image to the left is enhanced to increase visibility of viruses over a wide range of intensities. (E) Smaller viruses (<∼300nm) exhibit more variability in HA:NA ratio, which is bimodally distribution in viruses shorter than ∼300nm in length (population labeled i) but has an increasingly narrow distribution as virus size increases (populations from ii and iii).

Given the substantial variation we observe in the composition of individual virions, we next characterized covariation in the abundance of specific proteins. For this analysis, we focused on HA and NA, surface proteins with competing activities whose functional balance contributes to viral fitness (Wagner et al., 2002), and which represent, in the case of HA, a major target of vaccines and the adaptive immune response (Martinez et al., 2009; Sui et al., 2009), as well as the major source of antigenic drift (Koel et al., 2013). Variation in HA and NA abundance could significantly influence the binding characteristics of an individual virus particle (Xu and Shaw, 2016) and its ability to navigate environments rich in sialic acid, the ligand of HA and the substrate of NA. To determine the variation in relative abundance of HA and NA across a population, we measured the ratio between fluorescently-tagged HA and NA in individual particles. In contrast to the monomodal distributions of HA and NA measured individually, the ratio HA/NA is distributed bimodally in smaller (< ∼300 nm) virions and gradually reduces to a monomodal distribution as virus size increases from sub-diffraction limited spots to particles approaching ∼1µm in length (Figure 2D). Although viruses higher in NA are more likely to possess a portion of the viral genome, both subsets of the population contain a substantial fraction of particles that label positive for NP and thus are at least potentially infectious (63% out of 79004 higher-NA particles, versus 55% of 162174 lower-NA particles; Figure S7B).

### Compositionally and morphologically variable viruses are produced by single cells during one replication cycle

We next investigated the source of the compositional and morphological variability we observed. We reasoned that phenotypic variability in a population of viruses could arise from three general sources which are not mutually exclusive: (i) genetic diversity in the virus population, (ii) heterogeneity within the population of infected cells, or (iii) variability that arises as the virus is assembling. If phenotypic variability within a virus population requires genetic diversity or heterogeneity among infected cells, then a single virus infecting a single cell should not be capable of producing progeny that reflect the full diversity in morphology and composition seen in virus released from a monolayer of cells. To test this, we characterized the composition and morphology of progeny virus produced by individual cells infected with individual infectious units (MOI ∼ 0.01) over a single round of replication. To further reduce the likelihood of mutations arising during the replication cycle, we used a tagged virus strain that also contains a V43I mutation in PB1 that reduces the mutation rate by ∼1-3-fold (Cheung et al., 2014). To trap virus released by individual cells, we grew monolayers of MDCK epithelial cells in custom chambers consisting of a thick (∼4mm) collagen gel set within an acrylic enclosure whose height is offset above the collagen by ∼100µm (Figure 3A). Inverting these chambers and submerging in media above coverslips coated with anti-HA antibody positions the apical surface of the epithelial monolayer <100µm above the functionalized surface. Viruses budding from the apical surface of a cell in this orientation underwent limited lateral diffusion before becoming captured on the coverslip as they sedimented, creating clusters of immobilized viruses that originate from individual cells infected at low MOI. When we measure the composition of viruses located within separate clusters arising from this configuration, we observe similar distributions of HA-NA content and length as for virus raised in populations of cells (Figure 3B & C, Figure S8). This supports the hypothesis that the virus assembly process itself is ‘low fidelity’ and a primary contributor to phenotypic variability, and that neither genetic diversity in the virus population nor phenotypic variability among infected cells is needed for influenza A virus progeny to access a broad phenotypic space.

**Figure 3:**
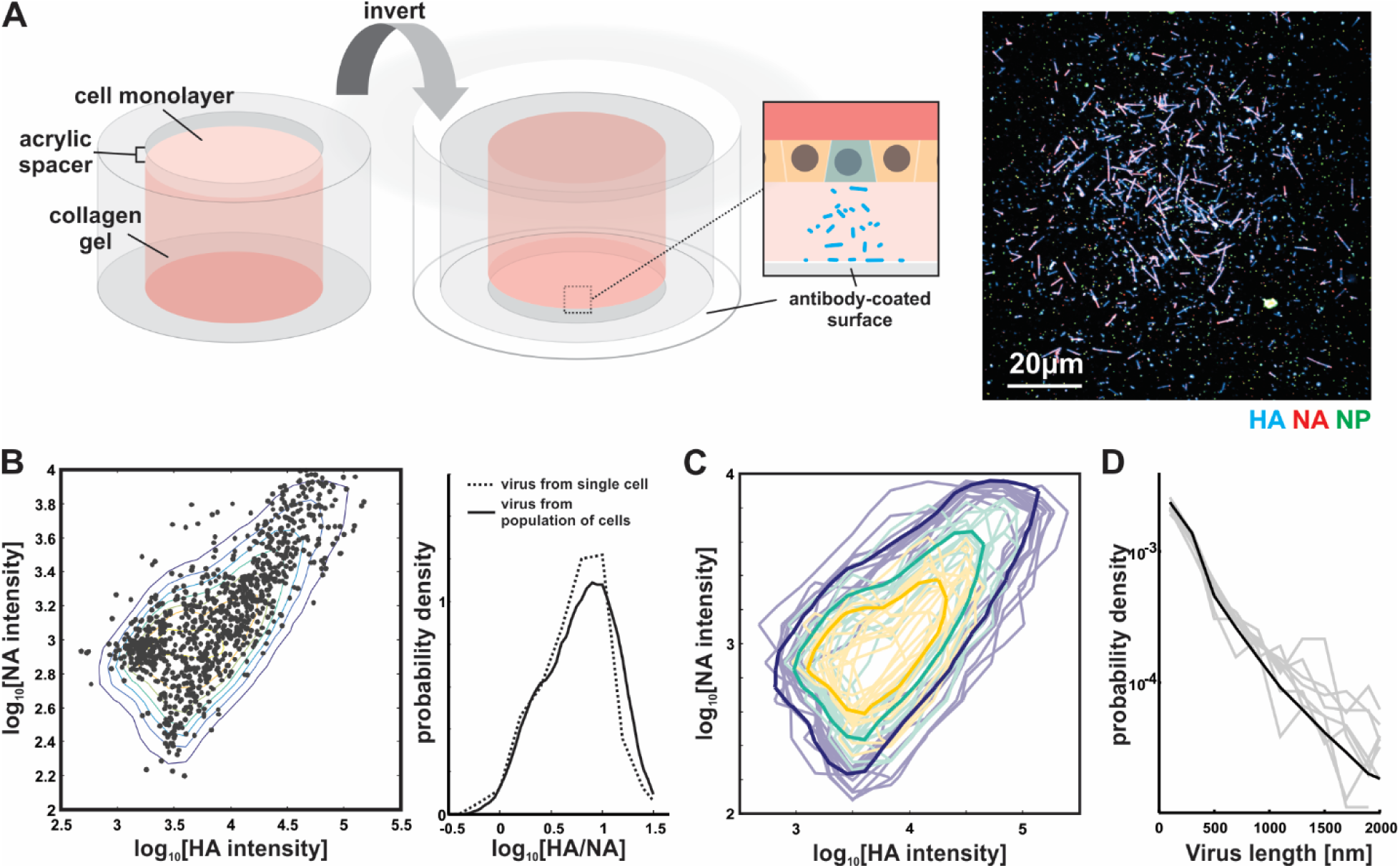
Phenotypic variability is encoded in the influenza A virus assembly process. (A) A method for characterizing viruses shed from individual infected cells. Viruses shed from an individual infected cell can be collected by positioning the apical surface of an epithelial monolayer tens of microns from a functionalized coverslip and infecting at low (∼0.01) MOI. (B) Groups of viruses shed from a single cell (shown as grey marks) exhibit similar compositional distributions (in both log-abundance of HA and NA, as well as the HA-NA ratio) as virus shed from a population of infected cells (shown as colored contours). (C) Overlayed distributions of virus shed from 12 separate clusters (average of 755 viruses per cluster) for HA-NA content, with light shading indicating data from a single cell and darker shading showing data across the population. (D) Overlayed length distributions from the same virus samples whose composition is plotted in (C).

To test whether phenotypic variability is influenced by environmental conditions during assembly in the absence of genetic differences, we changed the growth temperature of cells infected with virus bearing tagged HA, NA, and NP, along with the V43I mutation to PB1, from 37°C to 33°C (temperatures roughly representative of the lower and upper airways), after infecting for one hour at 37°C to control for potential temperature dependences in virus entry. After collecting virus at 16 hpi and evaluating virion composition through fluorescent labeling and imaging (Figure 4A & B), we found that lower temperature shifts the virus population towards higher NA content, even within a single round of replication, and that this shift is reversible from one generation to the next (Figure 4B). This plasticity in viral composition does not extend to all viral proteins, as the percentage of particles containing NP is not significantly affected by temperature shifts within this range (Figure 4C). This data provides further evidence that virus heterogeneity is not inherently linked to viral genotype and that aspects of viral phenotype – in this case, packaging of HA and NA, but not NP - can reversibly shift in the absence of sufficient time for mutations to accumulate.

**Figure 4:**
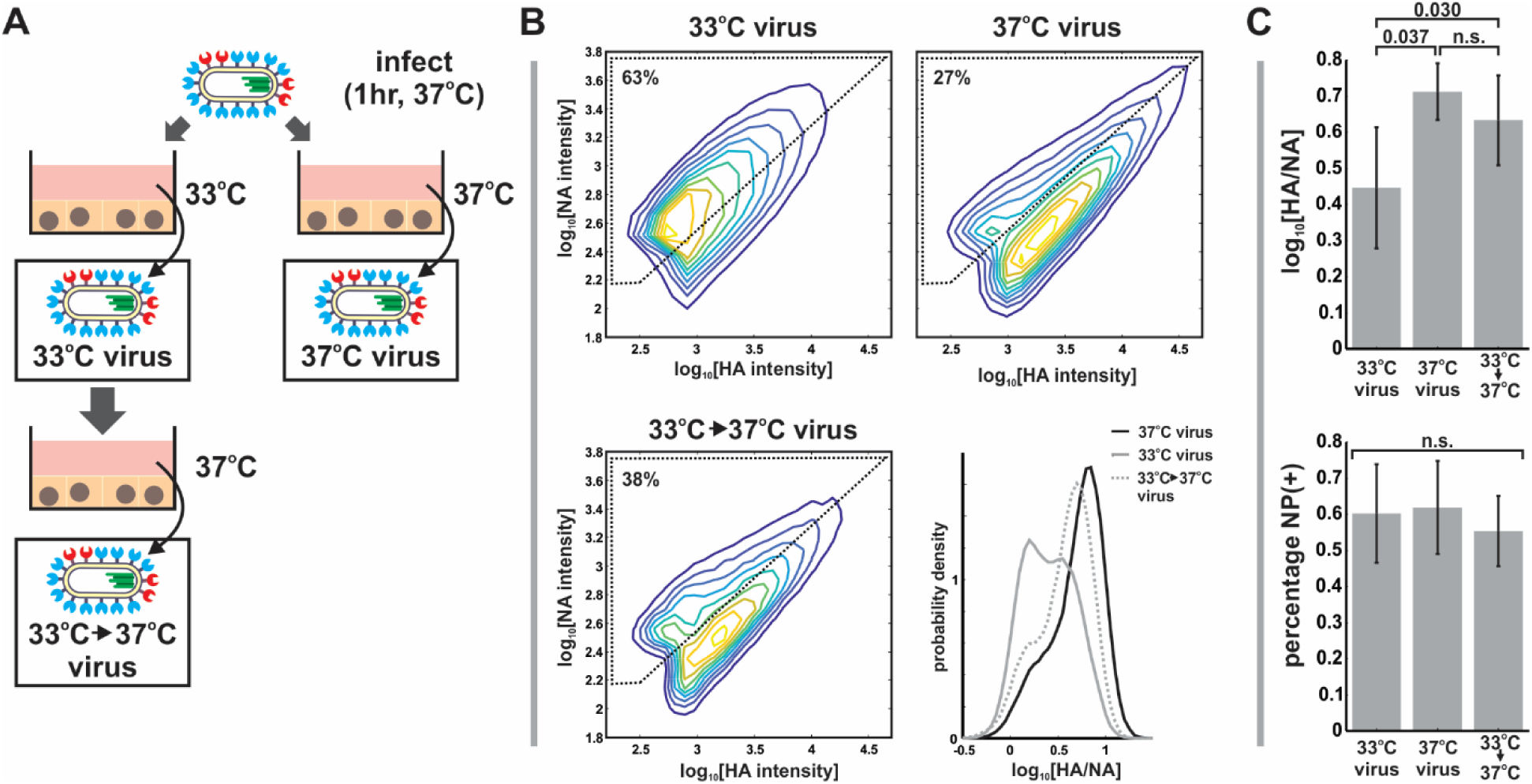
Compositional variability in individual viruses responds reversibly to changes in growth conditions. (A) Compositional variability in viruses raised under different environmental conditions was investigated by infecting cells at 37°C for one hour and allowing infection to proceed at either 33°C or 37°C. Viruses grown at 33°C for one generation are then returned to 37°C to evaluate reversibility of observed viral phenotypes. (B) Virus phenotype is malleable to environmental conditions, shifting to higher NA as the temperature is decreased from 37°C to 33°C. These shifts are reversible from one generation to the next, suggesting that they originate from phenotypic variability rather than changes in genotype. Plots show combined data from three biological replicates, with *N* > 30000 viruses per condition in each replicate. (C) In contract to the relative abundance of HA and NA (top), the fraction of particles labeling positive for NP is not significantly affected by temperature shifts between 33°C and 37°C. Quantification in all cases shows the mean and standard deviation from three biological replicates, with *N* > 30000 viruses per condition in each replicate. Significance evaluated using a paired-sample T-test.

### A subset of IAV populations escape challenge with NA inhibitors

The phenotypic variability of IAV allows individual virions to sample from a broad range of possible morphologies and compositions, resulting in a population of viruses with differing propensities to bind and cleave host receptors. This diversity could have an adaptive advantage in situations where the density of sialic acid varies unpredictably, as would occur during challenge with neuraminidase inhibitors (NAIs). To test how virus populations are shaped over short times by such a challenge (i.e. in the absence of genetic adaptations accumulating over multiple replication cycles), we characterized how treatment with a commonly used NAI – oseltamivir – influenced the phenotype of clonal, non-resistant virus populations within a single round of replication. To circumvent genetic adaptation to the NAI, we again used strains with the V43I mutation in PB1, and limited replication to a single round by infecting cells at MOI ∼ 1 and omitting trypsin from the growth media (Figure 5A). As NAI concentration was increased up to 1µM (an approximate upper bound for bioavailability in the sinuses (He et al., 1999; Kurowski et al., 2004)), released viruses become increasingly enriched in higher-NA virions and depleted in the higher-HA subset of the population (Figure 5B). For comparison, viruses that remained bound to the surface of infected cells under these same conditions were removed using an exogenous, oseltamivir-resistant sialidase (NanI from *C. perfringes*) and found to exhibit higher HA content and be more filamentous than viruses released from infected cells in the absence of sialidase treatment (Figure 5B, lower panels). These shifts are also apparent in the HA/NA ratio, which is significantly decreased (relative to the untreated sample) in viruses challenged with 100nM NAI and released from infected cells and significantly increased (relative to the treated sample) in viruses under the same conditions that remain bound to infected cells (and are released with exogenous NA) (Figure 5C).

**Figure 5:**
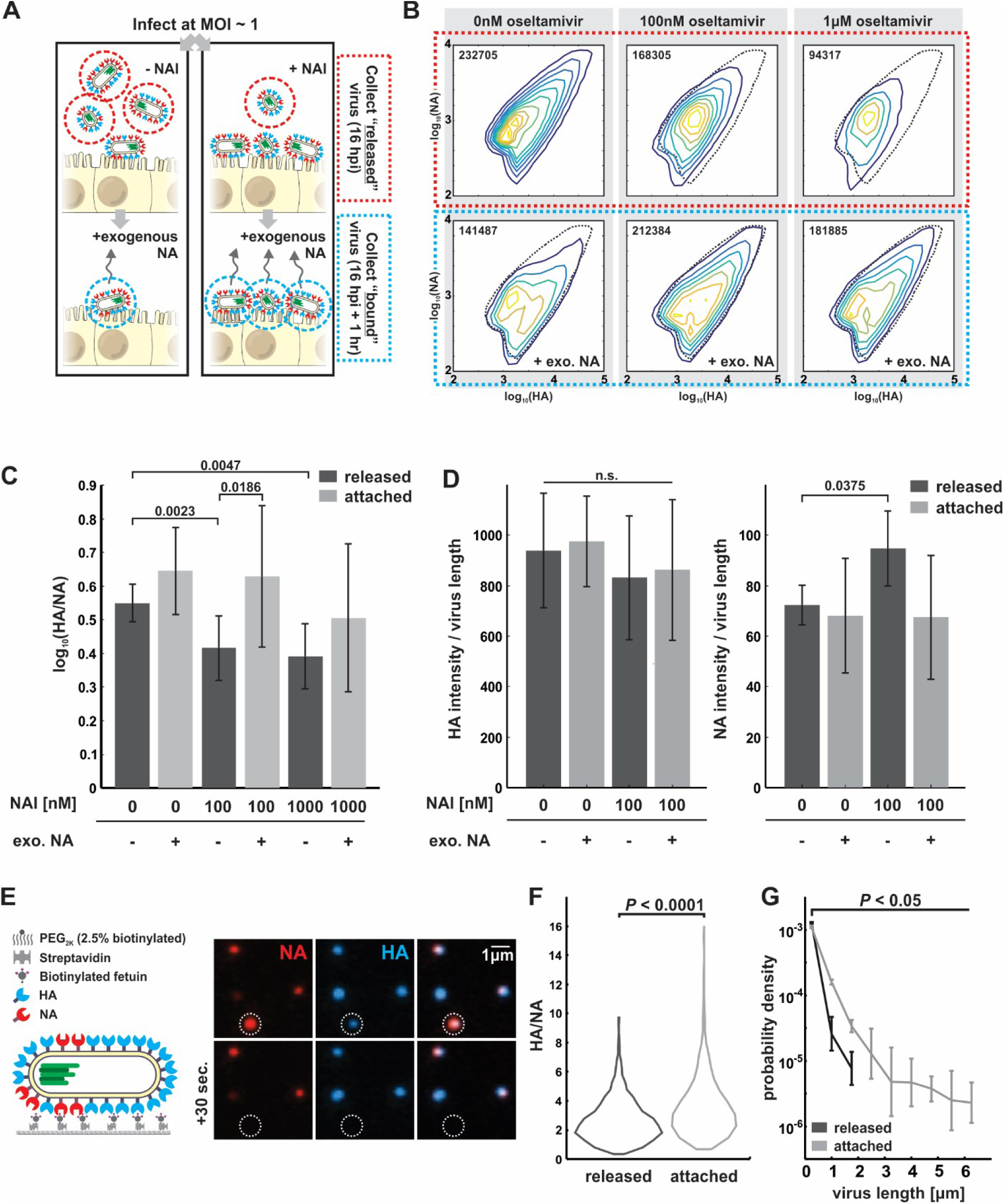
A compositional and morphological subpopulation of viruses challenged by neuraminidase inhibitors succeed in escaping from host cells. (A) To compare the heterogeneity of populations of virus exposed to NAIs that are able to detach from infected cells (red dotted circle) to those that are unable to detach by themselves (blue dotted circle), we first harvest virus from media at 16hpi, followed by the addition of media supplemented with an exogenous, oseltamivir-resistant bacterial neuraminidase. After one hour of treatment with exogenous NA, we harvest virus that had previously remained bound to the cell surface. (B) Treatment with physiological doses of the NAI oseltamivir shifts the phenotype of viruses that detach from the cell surface towards lower HA and higher NA abundance (top row). Viruses that remain attached to the cell surface under these conditions have predominantly high-HA phenotypes (“+exo. NA”, bottom row). (C) Comparison of HA-NA ratio in viruses released under different challenge (“NAI”) or permissive (“exo. NA”) conditions. Viruses that release from cells treated with 100nM or 1µM oseltamivir have significantly lower HA/NA ratios than those that release in the absence of inhibitor or in the presence of both inhibitor and exogenous NA. Bars represent median values +/-standard deviations for populations measured in at least three independent experiments per condition with a minimum of 94000 viruses each (*p*-values determined using a two-sample T-test). (D) Composition of filamentous viruses (*L* > 1µm) released under challenge and/or permissive conditions. While the density of HA on filamentous particles (defined as the total HA intensity divided by the particle length) does not change, filamentous viruses shed from cells challenged with NAI show a slight but significant increase in NA density. Bars represent median values +/-standard deviations for populations measured in four independent experiments with between 499 and 2067 filamentous viruses each (*p*-values determined using a two-sample T-test). (E) Measuring the propensity of viruses to detach from sialic acid covered surfaces as a function of their shape and composition. Coverslips modified with sialic acid at a density of ∼1 residue/nm^2^ provide a substrate for virus binding and release. (F) Viruses that detach within a 1hr observation period have significantly lower HA/NA ratios. (G) Viruses that detach within a 1hr observation period also have significantly shorter lengths. Quantification is based on three independent experiments with *N* > 366 released viruses and *N* > 991 attached viruses recorded in each. P-value for comparing HA/NA in released and attached populations calculated using a two-sample K-S test. P-value for comparing length distributions calculated using a two-sample T-test.

Shifts in the abundance of HA and NA in individual viruses could result from changes in virus morphology (as smaller viruses will have fewer of both proteins) as well as from changes in protein density on the viral surface. Although the surface density of receptor-binding HA on filamentous viruses (length > 1µm) does not differ significantly with NAI challenge, the average density of receptor-cleaving NA increases ∼30% after NAI challenge (Figure 5D), suggesting that packaging more NA confers an advantage in these conditions. This result indicates that HA/NA ratio contributes to the effective binding affinity of a virus. To test this idea more directly, we developed an *in vitro* virus detachment assay, in which we monitor the dissociation of viruses from sialic acid coated coverslips as a function of their morphology and composition (Figure 5E). For viruses bound to sialic acid at densities approximating the cell surface during NAI treatment (∼1 molecule/nm^2^ (Rosenberg, 1972)), those that dissociate over the course of an hour have significantly lower HA/NA ratios, consistent with the live cell experiments (Figure 5F).

The compositional balance between HA and NA has a clear impact on virus dissociation, but what about virus morphology? We find that morphology has a dramatic effect on detachment, with viruses that dissociate from the surface in our in vitro assay tending to be significantly shorter than those that remain stably bound (Figure 5G). Consistent with this, we observe substantial changes in the size distributions of released viruses under NAI challenge, with virion lengths >1µm significantly depleted from the pool of released virus (Figure 6A, left column). Although filamentous particles are recovered from NAI-treated cells following the addition of the exogenous sialidase (Figure 6A, right column), the total population of viruses produced in the presence of NAI treatment (i.e. those released naturally as well as those detached by sialidase treatment) reveals an overall reduction in virus length (Figure 6B). These data show that IAV can reliably escape from host cells treated with even high concentrations of NAI, but that the viruses released are only a subset of the phenotypically diverse population produced, with shorter virions having higher NA and lower HA better suited to survive NAI treatment. In addition to being filtered during NAI challenge, some phenotypic characteristics – in particular, virus morphology – are also malleable to environmental conditions, as illustrated by the shorter length distribution seen in the total virus produced during NAI challenge. However, other features of virus assembly such as the percentage of particles containing a genome (Figure S9) appear to be unaffected by NAI treatment. Taken together, the low fidelity assembly process of IAV appears to help virus subpopulations survive environmental fluctuations, including acute stresses, and quickly reestablish phenotypic diversity, all in the absence of genetic changes (Figure 7).

**Figure 6:**
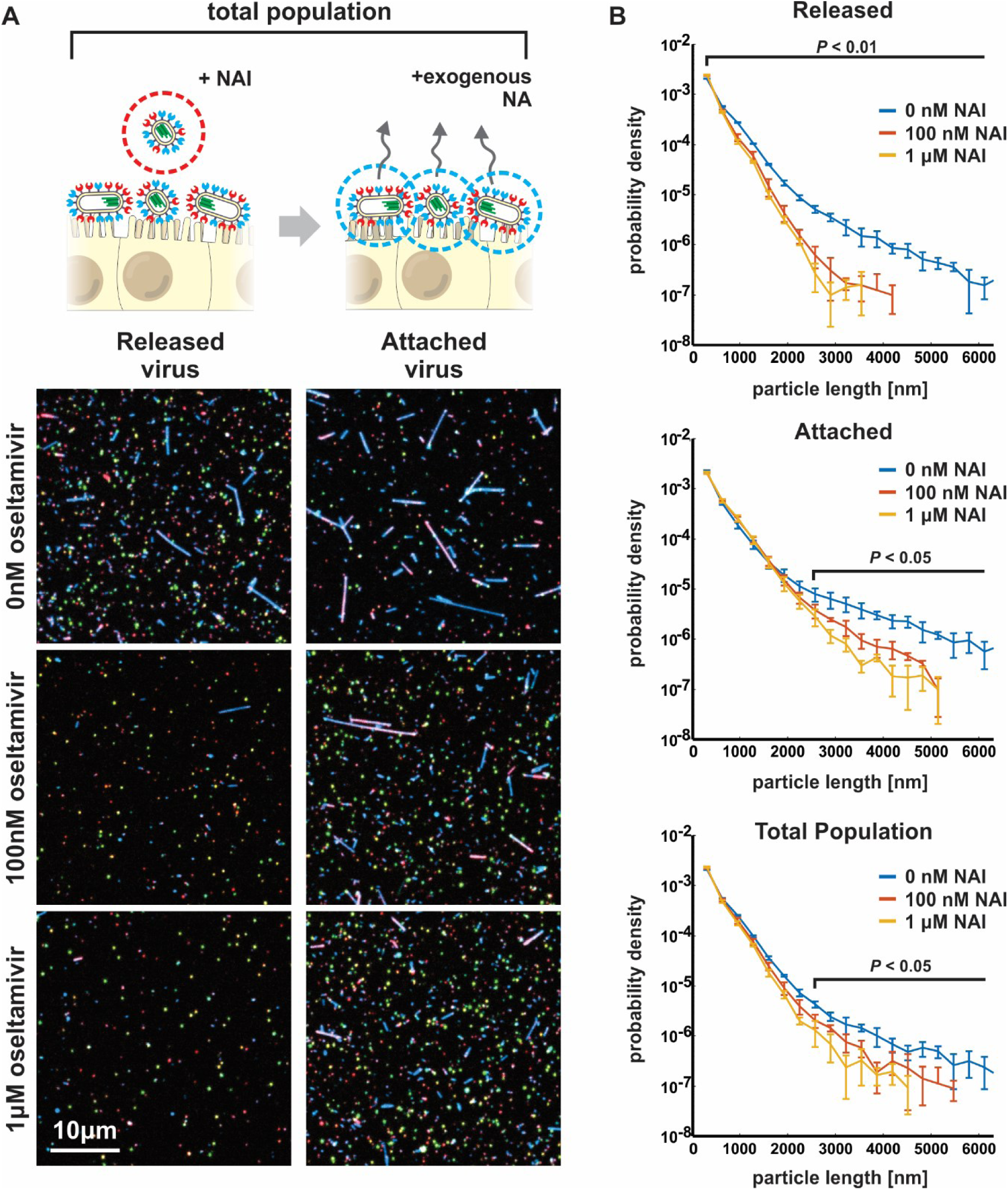
Morphological variability of influenza A virus is reduced by neuraminidase inhibitors challenge. (A) Treatment with NAI at approximate physiological concentrations shortens the typical length of released viral filaments (“Released” column), while longer filaments are preferentially sequestered on the cell surface (“Attached” column). (B) Frequency plots showing the distribution of particle lengths within released, attached, and total populations of viruses. Plots show the mean +/-standard deviation for length distributions determined from four independent experiments. For released viruses, differences are significant for all lengths (*P* < 0.01; two-sample t-test); for attached viruses as well as the total population, differences are significant for lengths above ∼2.5µm (*P* < 0.05; two-sample t-test).

**Figure 7:**
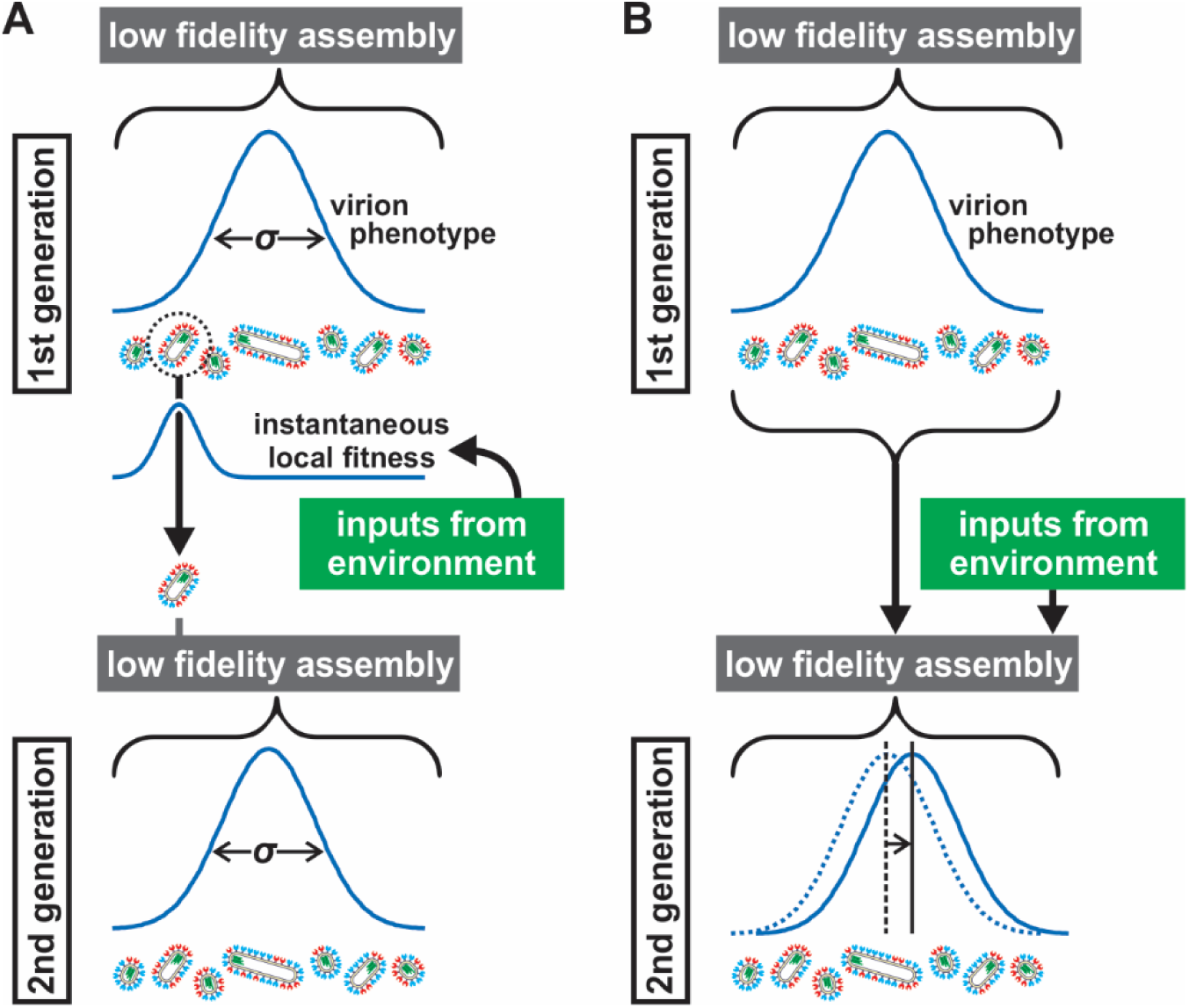
Low fidelity assembly in IAV contributes to survival in fluctuating environments by creating broad and malleable phenotypic diversity. (A) Phenotypic diversity within a population of influenza viruses (schematized by a distribution with variance *σ*) confers non-genetic fitness benefits on some subsets of the population that depend on environmental conditions. The low fidelity assembly of IAV allows these subpopulations (illustrated by a single surviving virus) to rapidly reestablish phenotypic diversity in the subsequent generation. (B) In addition to contributing to broad phenotypic diversity, IAV assembly also allows characteristics of a populations to reversibly shift in response to changing environmental conditions.

## Discussion

The infectiousness of a virus is a complex characteristic, arising from the specific activities of the viral proteins and the multiple barriers presented by the host. Success can often be predicted from genotype, which determines the cells to which the virus can bind and enter (Schrauwen and Fouchier, 2014), from the virus’s ability to suppress the innate immune response (Kochs et al., 2007), and from its ability to co-opt host factors to favor viral replication (Tripathi et al., 2015). As a result, mutations that alter these activities can have dramatic effects on host range and virulence, ultimately driving viral evolution. However, even as evolutionary processes shape viral populations over multiple generations, individual viruses face immediate and local challenges to productive infection and spreading. After budding from an infected cell, the virus must dissociate from the cell surface, navigate the mechanical and chemical barriers imposed by mucociliary clearance, avoid neutralization from innate and adaptive immunity, and bind to a target cell in a manner that enables internalization and subsequent fusion between virus and host membrane. The ability of an individual virion to accomplish these tasks depends both on the genetically-specified activity of its proteins as well as factors that may not be under direct genetic control: the shape of the virus and the abundance of viral and cellular proteins that are packaged during assembly. In the case of influenza A virus, a highly pleiomorphic virus, significant variability in morphology and composition occurs within the population. Because the contributions of these factors can be genetically silent, their significance has been difficult to determine and often overlooked in laboratory studies of infectivity. However, the long-standing observation that *in vivo* replication favors forms of influenza A virus that have more heterogeneous morphology – a characteristic that is genetically influenced by IAV’s matrix protein M1 – suggests that phenotypic variability may confer benefits to IAV under certain circumstances.

To better understand how virus phenotype contributes to the replicative fitness of influenza A virus, we developed strains of IAV that harbor tags on each of the virus’s main structural proteins (Figure 1). This enabled us to characterize phenotypic variability in influenza A virus populations by measuring virus size and the abundance of specific viral proteins (Figure 2). In the present work, we have focused on HA and NA, the most abundant proteins in the viral membrane and competitors for binding and destruction of sialic acid, the virus receptor. Our findings demonstrate that heterogeneity in abundance of HA trimers and NA tetramers at the individual virus level changes the balance of intrinsic protein activity, creating virions with varying proclivity towards receptor binding or receptor destruction. Even in a genetically uniform population, the low fidelity assembly process of influenza A virus that samples broadly from the possible distributions of HA, NA, and size gives rise to wide phenotypic variability that results in functionally relevant differences in virus binding and release.

Why might influenza A virus favor heterogeneous progeny? One possible answer is that phenotypic variability increases the probability of survival in an unpredictable environment (Kussell, 2005). Creating diverse progeny could contribute to the persistence of a virus population and allow sufficient time for an adaptive genetic response to evolve. Phenotypic adaptation would be particularly valuable during a population bottleneck when genetic diversity is limited, such as during the initial infection of a new host (Poon et al., 2016; Varble et al., 2014). Consistent with this concept, we find that the phenotypic variability of viruses released from individual cells infected with individual infectious units is comparable to that produced by populations of cells infected with populations of viruses (Figure 3), indicating that a single virus can rebuild phenotypic diversity in a single round of replication. A second feature of phenotypic variability that compliments genetic diversity is its non-committal nature: a characteristic that is favorable in one circumstance may be detrimental in another. If a single virus can generate a phenotypically diverse population and only one member of that population needs to survive for replication to continue, maintaining that diversity across generations would offer an adaptive advantage. In support of this, we show that IAV phenotype can vary reversibly (i.e. viruses with higher NA content are not genetically predisposed to make progeny with higher NA content, Figure 4), demonstrating that adaptation through variability in composition or morphology does not constrain the infectiousness of subsequent generations. In this way, phenotypic variability could serve as a buffer against environmental fluctuations – including those induced by treatments with antiviral agents (Figures 5 & 6).

The benefits of phenotypic variability discussed here would not be exclusive to IAV, raising the possibility that similar strategies may be harnessed by other viruses. Interestingly, pleomorphism is common among enveloped viruses, with filamentous morphology common in the filoviridae, pneumoviridae, and paramyxoviridae families, in addition to the example of IAV within orthomyxoviridae (Compans et al., 1966; Kuhn et al., 2010; Roberts et al., 1995). Even within virus families where particle morphology is comparatively uniform (e.g. flaviviridae), other forms of functionally-relevant phenotypic variability may be present, such as in the balance between immature and mature glycoproteins on the viral surface (Pierson and Diamond, 2012). More generally, the size, composition, and epitopes displayed on the surface of many viruses influence how they gain entry into a cell and interact with the innate and adaptive immune response (Dejnirattisai et al., 2016; Geng et al., 2007; Rossman et al., 2012), suggesting that variability in these properties could offer fitness advantages. Even defective viral particles that are not themselves infectious can enhance survival and persistence of the infectious virions produced alongside them (Xu et al., 2017). Given that a single, optimal path to infection likely does not exist, the ability to produce diverse progeny without making an evolutionary commitment to one particular path – and one stereotyped virion – may contribute to the fitness of a wide variety of viruses.

The underlying mechanisms responsible for IAV’s phenotypic variability remain an active topic of research. While our understanding of the mechanisms by which composition and morphology in IAV is established remains incomplete, there is emerging consensus that the assembly process involves contributions from many components, with some functional redundancy likely (Ali et al., 2000; Chlanda et al., 2015; Rossman and Lamb, 2011). The ability of IAV to assemble through multiple pathways could contribute to the low fidelity assembly we observe within populations of progeny virions. Analogous to protein folding and other systems involving complex networks of interactions, different stages in the assembly of a virus could in principle be either thermodynamically controlled or kinetically controlled (Baker and Agard, 1994; Cui et al., 2007). As demonstrated by in vitro reconstitution experiments and computational modeling, highly-ordered viral capsids can, under certain circumstances, successfully avoid kinetic trapping to reach a uniform, minimum energy state (Butler and Klug, 1971; Hagan and Chandler, 2006; Zlotnick et al., 2000). However, we propose that influenza A virions may be kinetically trapped structures, with HA and NA content locked in during assembly through interactions with other structural proteins and unable to equilibrate before the reaction is terminated by membrane scission. In this view (which is consistent with the shifts in virus populations we see at different growth temperatures), stochastic incorporation of proteins (HA, NA, and host proteins) from the local plasma membrane into the viral envelope could account for compositional variability, while morphological variability could arise from stochasticity in the timing or mechanism of membrane scission.

Our findings provide a first assessment of phenotypic variability at the individual virion level in live influenza A virus populations and offer evidence that this variability can promote virus survival. However, these experiments focus on a single laboratory strain of IAV engineered to produce filamentous particles *in vitro*. More work is needed to determine the generality of these findings to different strains and subtypes, as well as to *in vivo* models of infection. The tools we describe here for minimally disruptive fluorescent labeling of influenza A virus provide a means of assessing this generality and a conceptual guide for asking similar questions about other filamentous viruses. The large variability that we measure in HA/NA ratio raises the possibility that other aspects of virus composition and organization may be similarly heterogeneous. In particular, the abundance of other viral and cellular proteins and lipids packaged during assembly could have consequences on the rate of acidification of the virus interior (Ivanovic et al., 2012), the route the virus follows through the endosome (He et al., 2013), the uncoating of the virus following fusion (Banerjee et al., 2014), and the general chemical and physical properties of the viral membrane (Gerl et al., 2012; Polozov et al., 2008). Since adaptive and therapeutic interventions against IAV could alter viral phenotypes in genetically-independent ways, direct characterization of viral populations in the manner described here may help in evaluating the impact of emerging antiviral therapies (Furuta et al., 2013; Kadam et al., 2017; Strauch et al., 2017). Overall, understanding the extent to which viral heterogeneity contributes to viral persistence could help to guide the development of a new class of therapies aimed at reducing natural variability, or applying combinatorial pressure that mitigates the survival benefits of phenotypic variability.

## Acknowledgements

The authors would like to thank Fletcher Lab members for helpful feedback and technical consultation. This work was supported by the NIH R01 GM114671 (DAF), the Immunotherapeutics and Vaccine Research Initiative at UC Berkeley (DAF), and the Chan Zuckerberg Biohub (DAF). M.D.V. was funded by a Burroughs Wellcome Fund CASI Fellowship. D.A.F. is a Chan Zuckerberg Biohub investigator.

